# 3D Graph Contrastive Learning for Molecular Property Prediction

**DOI:** 10.1101/2022.12.11.520009

**Authors:** Kisung Moon, Hyeon-Jin Im, Sunyoung Kwon

## Abstract

**Motivation:** Self-supervised learning (SSL) is a method that learns the data representation by utilizing supervision inherent in the data. This learning method is in the spotlight in the drug field, lacking annotated data due to time-consuming and expensive experiments. SSL using enormous unlabeled data has shown excellent performance for molecular property prediction, but a few issues exist. (1) Existing SSL models are large-scale; there is a limitation to implementing SSL where the computing resource is insufficient. (2) In most cases, they do not utilize 3D structural information for molecular representation learning. The activity of a drug is closely related to the structure of the drug molecule. Nevertheless, most current models do not use 3D information or use it partially. (3) Previous models that apply contrastive learning to molecules use the augmentation of permuting atoms and bonds. Therefore, molecules having different characteristics can be in the same positive samples. We propose a novel contrastive learning framework, small-scale 3D Graph Contrastive Learning (3DGCL) for molecular property prediction, to solve the above problems.

**Results:** 3DGCL learns the molecular representation by reflecting the molecule’s structure through the pre-training process that does not change the semantics of the drug. Using only 1,128 samples for pre-train data and 0.5 million model parameters, we achieved state-of-the-art or comparable performance in six benchmark datasets. Extensive experiments demonstrate that 3D structural information based on chemical knowledge is essential to molecular representation learning for property prediction.

**Availability:** Data and codes are available in https://github.com/moonkisung/3DGCL.

## 1 Introduction

Self-supervised learning (SSL) learns data semantics by utilizing inherent supervision in data from a large amount of unlabeled data. A self-supervised model using a pretext task has recently outperformed the general supervised model. SSL has been extremely successful in computer vision (Chen *et al*., 2020) and language fields (Mikolov *et al*., 2013; Devlin *et al*., 2018) and has attracted particular attention in the field of drugs (Rong *et al*., 2020), wherein considerable time and money are incurred labeling data, and there is a lack of annotated data compared with other domains. A molecule can be expressed in various ways, such as a chemical fingerprint, for example, ECFP (Rogers and Hahn, 2010), which uses a fixed vector for particular substructures, and simplified molecular input line entry system (SMILES), (Weininger, 1988) which represents the molecule as a string.

In addition, there is a way to represent a molecule as a graph, and Graph Neural Networks is widely used for molecular property prediction (Gilmer *et al*., 2017) because it can reflect the structure and correlation of atoms and bonds effectively. SSL using enormous amounts of unlabeled data has shown excellent performance for molecular property prediction (Li *et al*., 2021; Zhang *et al*., 2020). However, a few issues exist. First, Existing self-supervised learning models are ‘large-scale.’ They require a million sizes of pre-train data to generalize various downstream tasks and, in many cases, are large-size models such as Transformer (Vaswani *et al*., 2017) to learn that data. Therefore, there is a limitation to implementing self-supervised learning where the computing resource is insufficient. We use only 1,128 samples for pre-train data, about 0.5 million model parameters, and overcome the not-high computing environment. Second, in most cases, self-supervised models do not utilize 3D structural information for molecular representation learning. The activity and property of a drug are closely related to the structure of the drug molecule. Nevertheless, most current self-supervised models do not use 3D information or use it partially (Liu *et al*., 2021b; Stärk *et al*., 2021). We introduce a novel 3D-3D view contrastive learning method to learn molecular structural-semantic. Contrastive learning is one of the self-supervised learning methods and consists of pretext tasks to learn similarities and dissimilarities between positive and negative pairs. Finally, previous models that apply contrastive learning to molecules have used the augmentation permuting atoms and bonds, while positive samples should be intrinsically identical to each other. Unlike images, molecules can be completely different if we use the augmentation that changes atoms or bonds, so molecules having different characteristics can be in the same positive samples. We generate a conformer pool consisting of several conformers to preserve the molecular composition and use it for molecule-contrastive learning.

We present a 3D Graph Contrastive Learning (3DGCL) framework for molecular property prediction, a small-scale method that uses a tiny dataset, model, and 3D coordinates. Our approach uses approximately 1,000 data and 0.5 million model parameters, and randomly selects molecules from the conformer pool instead of selecting the most stable molecules to learn the 3D structure abundantly. We demonstrated the effectiveness of the 3DGCL through extensive experiments. We compared the proposed method with previous state-of-the-art baselines in four regression and two classification benchmark datasets under the same experimental settings. To investigate the importance of the 3D view, we compared our method of utilizing molecular 3D coordinates with the existing pre-training method of modifying the original molecule. In addition, we also compare our conformer pool, comprising conformers that exist in nature, with molecules that are difficult to exist in nature, which is created by adding noise to the structure of the original molecule. We achieved outstanding performance and the best result in the pre-training effect by comparing the difference between pre-training and non-pre-training, compared to existing methods. The results showed that chemical-based 3D structural information is vital for molecular representation learning in property prediction. We focus not only on large-scale learning but also on pre-training strategies to learn representations correctly in self-supervised learning.

Our main contributions are:

- We develop a compact self-supervised learning approach that can be run even in environments with low computational resources, using the small-scale pre-train samples and parameters. We also achieve the state-of-the-art or comparable performance in six molecular benchmarks.
- To the best of our knowledge, we propose 3D-3D view contrastive learning that can take full advantage of 3D information for the first time. We actively utilize 3D positional information inherent in molecules through a pre-training scheme using a conformer pool.
- Extensive experiments demonstrate that our method, which can utilize structural information abundantly while maintaining semantics, is more suitable for molecular property prediction than conventional contrastive learning, which may change the structure or properties of molecules.

## 2 Related Works

### 2.1 Graph Neural Networks

Graph Neural Networks (GNNs) is a widely used deep learning technique using graph-structured data. Due to the fact that molecules can be well described in graphs, GNNs for molecular property prediction has been active research (Yang *et al*., 2019a; Gilmer *et al*., 2017; Lu *et al*., 2019). The molecular graph is represented as 𝒢 = 𝒱, *ε*), where 𝒱 and *ε* denote the set of atoms and bonds, respectively.

The message passing scheme in GNNs (Gilmer *et al*., 2017) can be formalized as follows:

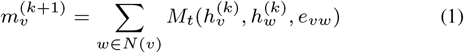

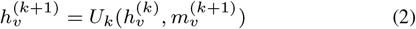

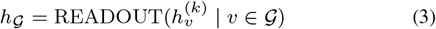

The GNNs aims to learn each node vector 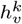 and the entire graph vector *h*_𝒢_ in the k-th layers. *N* (𝓋) denotes the neighbor node of node 𝓋 and *e*_𝓋𝓌_ denotes the edge between node 𝓋 and node 𝓌. *M* and *U* are message and update functions depending on the GNN models. GNNs update each node through an iterative message passing process. Finally, the readout layer, e.g. sum or mean pooling, is applied to get the entire graph vector while satisfying permutation-invariance.

### 2.2 Self-supervised learning for molecular property prediction

In the drug field, it is challenging to obtain annotated data due to expensive wet-lab experiments; however, it is relatively easy to collect data without annotation. Thus, there have been many SSL approaches recently, and they have shown noticeable results for molecular property prediction.

Inspired by BERT (Devlin *et al*., 2018), a powerful pre-training model in NLP, SMILES-BERT (Wang *et al*., 2019), ChemBERTa (Chithrananda *et al*., 2020) utilized the enormous SMILES datasets to predict molecular properties. However, SMILES is not geometry-aware because it represents molecules as string sequences. Instead of SMILES as molecular representations, recent studies using molecular graphs use various types of pretext tasks. GROVER (Rong *et al*., 2020) developed a transformer-style architecture and utilized motif-level pretext tasks. (Dillard, 2021) integrated pretext tasks at a different scale consisting of the atom, fragment, and molecule levels.

Contrastive learning is one of the powerful learning methods of self-supervised learning, making similar samples close and dissimilar samples far away in embedding space (He *et al*., 2020; Chen *et al*., 2020). GraphCL (You *et al*., 2020) proposed various augmentation methods for graph contrastive learning. MolCLR (Wang *et al*., 2022) focused on molecular representation learning based on approaches presented in GraphCL. These methods change the structure of the original graph, such as dropping nodes or changing edges, so that molecules can have different properties. Zhang *et al*. (2020) proposed contrastive method brings the subgraph and the graph close together through motif-level sampling from the entire graph. MoCL (Sun *et al*., 2021) used augmentation to replace the sub-structure in the original molecule with bioisostere, which has similar properties to the original, incorporating domain knowledge. They tried to maintain the semantic information of the molecule, but supervision used in pre-training may not be appropriate for molecular representation learning because even a tiny change in a molecule can lead to a significant difference. Hermosilla and Ropinski (2022) proposed a 3D protein contrastive learning method to learn the structure of a protein. During the pre-training process, the substructures of the protein are used, which is a similar approach to (Sun *et al*., 2021) except that 3D information is used. MEMO (Zhu *et al*., 2022) is a multiple-view contrastive learning method using various molecular representations (2D graph, 3D graph, Fingerprint, and SMILES). Uni-Mol (Zhou *et al*., 2022) is a self-supervised model that uses 3D information and consists of a pre-training scheme that predicts masked atoms or denoises 3D positions after adding noise to the molecular coordinates.

Molecular structure information plays an essential role in determining molecular properties. We have recently witnessed studies with great success using 3D geometric data. GraphMVP (Liu *et al*., 2021b) and 3Dinformax (Stärk *et al*., 2021). developed 2D-3D view contrastive learning approaches maximizing the mutual information between molecular embedding of 2D graph network and 3D graph network. Existing works with 3D coordinates do not fully utilize geometric information because they allow the 2D view network to learn 3D view information.

### 2.3 3D Graph Neural Networks

Due to a general graph being represented in a non-euclidean space, it is difficult to grasp the exact molecular structure. However, by leveraging 3D positions, we can specifically use unique molecular geometric properties such as distance, angle, and torsion in the euclidean space. For this reason, 3D Graph Neural Networks (3D GNNs) using 3D information are attracting widespread attention in drug and material discovery fields (Unke and Meuwly, 2019; Qiao *et al*., 2020).

3D Molecular graph can be represented as 𝒢^3𝒹^ = (𝒱, *ε*, ℛ), where ℛ denotes the 3D coordinates of atoms. SGCN (Danel *et al*., 2020) applied different weights according to interatomic distance in the message-passing process based on GCN. SchNet (Schütt *et al*., 2017) used Gaussian radial basis functions to represent distance information in a high-dimensional space. DimeNet (Klicpera *et al*., 2020) used orthogonal functions to learn both distance and angle information. SphereNet (Liu *et al*., 2021c) also used torsion information to represent a complete molecular structure using spherical message passing. ChIRo (Adams *et al*., 2021) developed a chirality-aware 3D network that can learn molecular chirality from torsion angles. SchNet and subsequent models (DimeNet and SphereNet) can capture non-bonded interaction based on 3D positional information within cutoff. SchNet is the most efficient architecture among 3D GNNs in terms of the learning cost and time, although the other models are superior performance. We use SchNet as an encoder for 3D contrastive learning. The critical process of SchNet is expressed as follows:

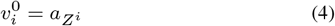

where 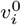 denotes the feature of the atom *i* and is the initialized value using embedding of the atomic number.

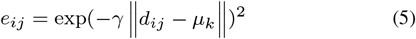

where *d*_*ij*_ denotes the distance between the atom *i* and the atom *j*. The interatomic distance is encoded to *e*_*ij*_ by radial basis functions located in k centers *μ*_*k*_.

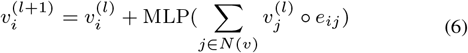

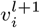 in the *l*+1-th layer is updated based on neighbor atom *j* in the *l*-th layer and the distance information *e*_*ij*_. ○ indicates the element-wise multiplication.

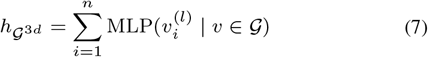

The global molecular feature vector 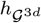 is obtained based on summation of the atom embeddings. We can handle arbitrary positioned atoms in the 3D space through the encoder.

## 3 Method

### 3.1 Conformer Pool

A conformer is a molecule group formed by rotation on single bonds in a molecule. Conformers have different potential energies depending on the degree of rotation, and the lower the energy, the higher the probability of existence in nature. For example, butane is a molecule composed of single carbon-carbon bonds and single carbon-hydrogen bonds. If it is expressed in 2D, it is difficult to see the change due to the rotation of a single bond. Nevertheless, as shown in Figrue 1, if represented in 3D, we can see that butanes form conformers with different potential energies according to rotation. We generate a conformer pool using the Merck molecular force field (MMFF94) (Halgren, 1996) function in RDkit (Landrum *et al*., 2020) to utilize diverse molecular geometric information in contrastive learning. The MMFF method combines distance geometry (DG) algorithms, a classic approach that randomly sample conformational space, with energy minimization using MMFF. The conformer pool consists of five conformers with the lowest energy. We do not add more conformers because five conformers are enough to represent almost all molecules in nature (Liu *et al*., 2021b), and the more conformers we add, the more computational cost we need. We also reduce the cost and enrich the molecular representation by randomly selecting the conformers from the conformer pool.

**Fig. 1.**
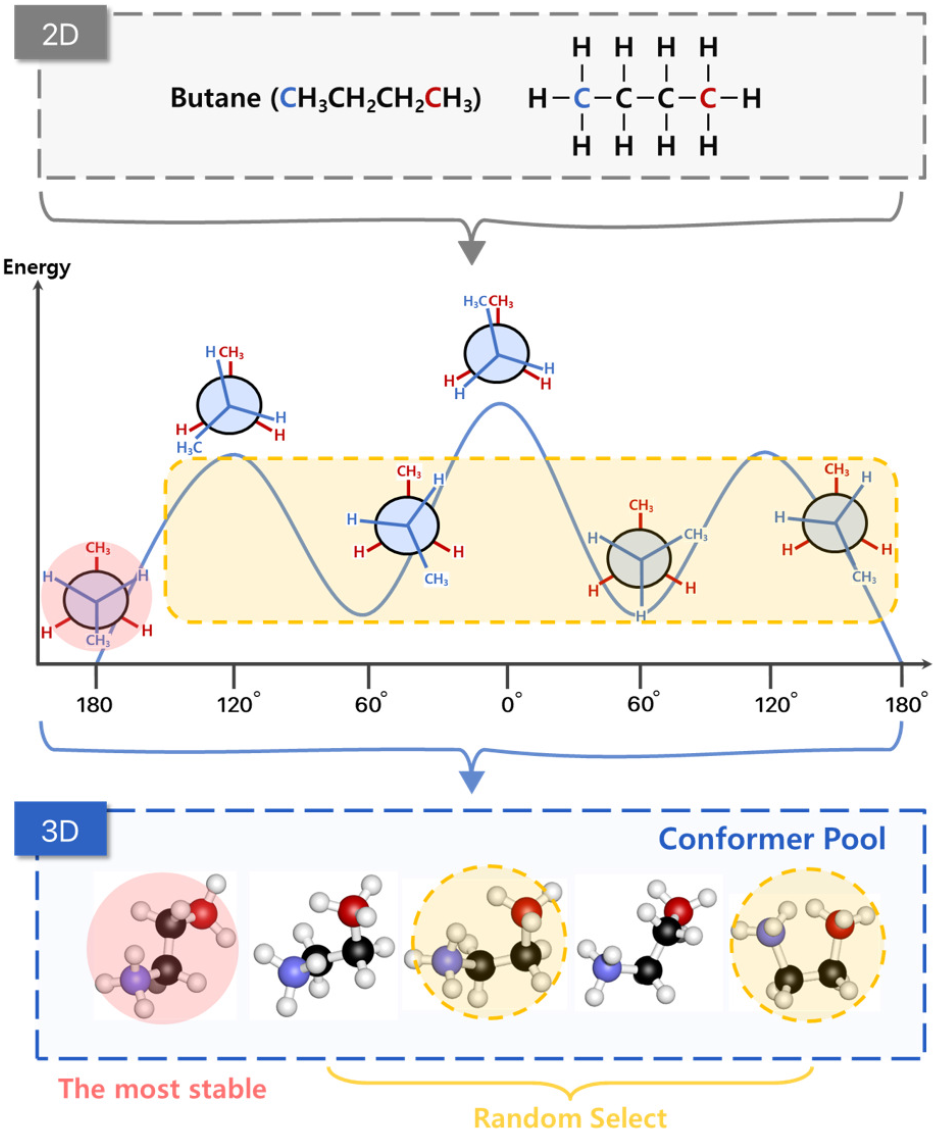
The process of creating a conformer pool we use in contrastive learning. We cannot grasp the spatial characteristics of butane in 2D. However, we can see that the potential energy varies depending on the rotation angle of the bond between the second carbon (C2) and the third carbon (C3), both colored black in the figure on top. We construct a pool of conformers with different stability and pre-train by randomly selecting from the rest of the conformers shown in the yellow box in the figure in the middle, except for the most stable molecule.

### 3.2 3D Graph Contrastive Learning

Contrastive learning is a powerful self-supervised learning method, moving positive pairs of similar samples close and negative pairs of dissimilar samples far away in embedding space. In order to be consistent with the basic assumption of contrastive learning, we make the positive samples be intrinsically the same as each other, unlike previous works that have made use of the augmentation permuting atoms and bonds or may not maintain molecular semantics (Wang *et al*., 2022; Zhang *et al*., 2020; Sun *et al*., 2021).

Maintaining semantics between conformers means that conformers conserve identity with the same element. Therefore, we performed contrastive learning by constructing positive pairs with conformers (molecules with slightly different properties) instead of the existing methods of changing semantics in the pre-training process.

Given molecular-input samples, we augment the molecule by applying our conformer pool to construct the positive and negative pairs. We randomly select two conformers (in yellow circle) in the conformer pool, as shown in Fig. 2.

**Fig. 2.**
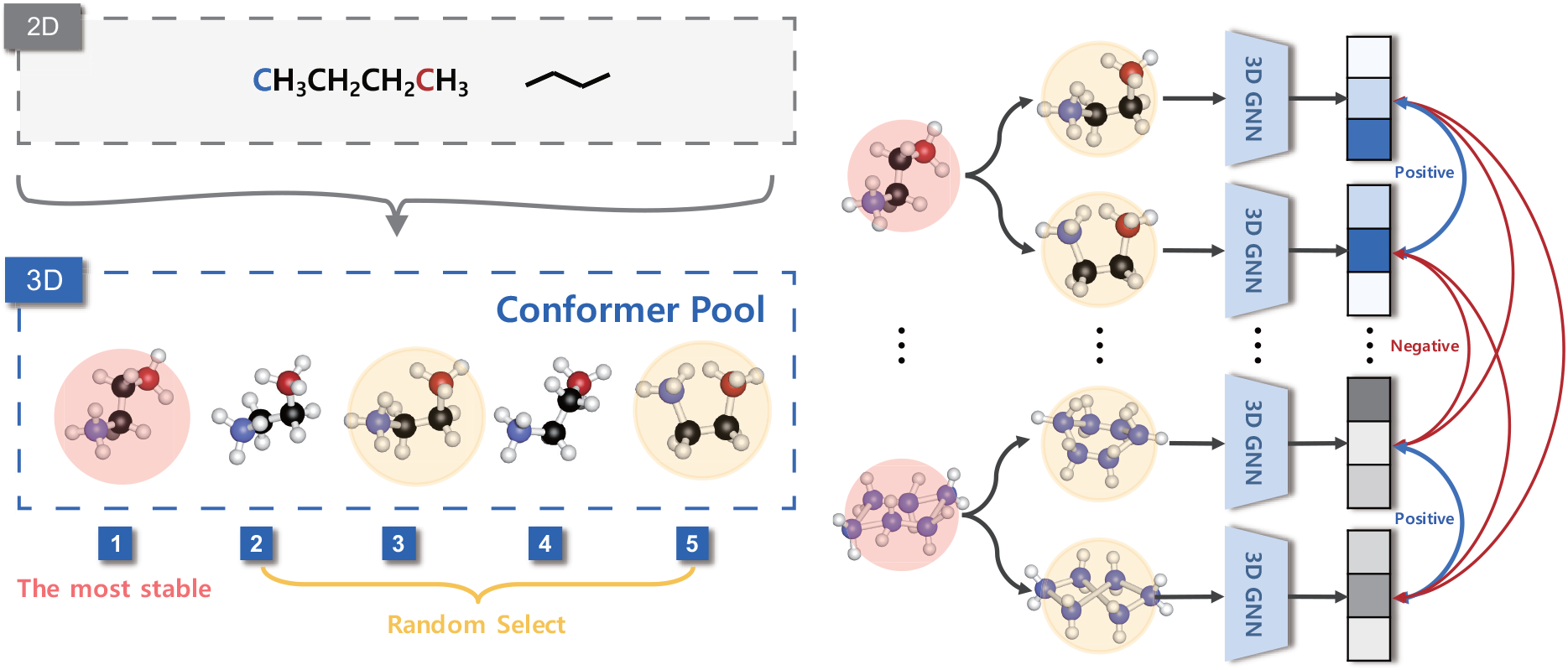
We generate a conformer pool from the original 2D molecule using 3D coordinates information. The conformer pool consists of five conformers that vary stability with the rotation angle of a single bond. We randomly choose the two molecules from the conformer pool for conformer augmentation. Then, the selected molecule is fed to a 3D encoder for embedding molecular representation. We apply a projection head to map the 3D view representations into the latent space for contrastive learning. The contrastive loss is computed across all positive and negative pairs in the minibatch of N molecules.

Graph contrastive learning maximizes the consistency between latent representations of positive pairs against negative pairs. We get the molecular graph embedding *h* using a 3DGNN encoder as the SchNet, and we apply a projection head to embedding *h* resulting in the non-linear transformation to obtain latent representation *z*. The projection head consists of a two-layer multi-layer perceptron and maps the molecular representations to another latent space as advocated in (Chen *et al*., 2020). Finally, we adopt a normalized temperature-scaled cross entropy (NT-Xent) (Chen *et al*., 2020) loss as our objective loss to maximize the agreement between the positive samples compared to the negative samples in a minibatch of N molecules. The contrastive loss for the first and second sampled conformers of k-th graph *k*_*c*1_, *k*_*c*2_ is defined as:

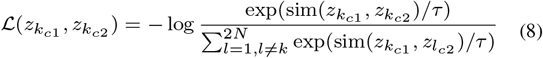

where sim (*z*_1_, *z*_2_) is the cosine similarity *z*_1_ *· z*_2_/(∥*z*_1_∥ ∥*z*_2_∥) and *τ* is the temperature paramerter. We compute the final loss across all training samples in the minibatch. There have been a few molecular graph contrastive learning models using diverse viewpoints recently. As introduced in the Section 2.2, MoCL and MolCLR (Sun *et al*., 2021; Wang *et al*., 2022) use the 2D-2D view approach, and there are models 3Dinformax and GraphMVP (Stärk *et al*., 2021; Liu *et al*., 2021b) that use the method of the 2D-3D view. However, as far as we know, we propose a 3D-3D perspective contrastive learning model first to fully utilize 3D information in the context of molecular property prediction, as shown in Table 1.

**Table 1.** Comparison of views in various contrastive learning models.

| Model | 2D-2D view | 2D-3D view | 3D-3D view |
| --- | --- | --- | --- |
| MoCL | ✓ | · | · |
| MolCLR | ✓ | · | · |
| 3Dinformax | · | ✓ | · |
| GraphMVP | · | ✓ | · |
| **3DGCL** | · | · | ✓ |

## 4 Experiments

### 4.1 Datasets

*Pre-training Dataset* For pre-training of 3DGCL. We use only 1,128 ESOL datasets (Delaney, 2004). We add 3D coordinates to pre-train datasets using the Merck molecular force field (MMFF94) (Halgren, 1996) function, which can obtain 3D coordinates faster (Stärk *et al*., 2021) than the latest deep learning-based methods (Ganea *et al*., 2021; Shi *et al*., 2021).

#### Downstream Datasets

We use four regression and two classification datasets for downstream tasks. The datasets are described in Table 2:

**Table 2.**
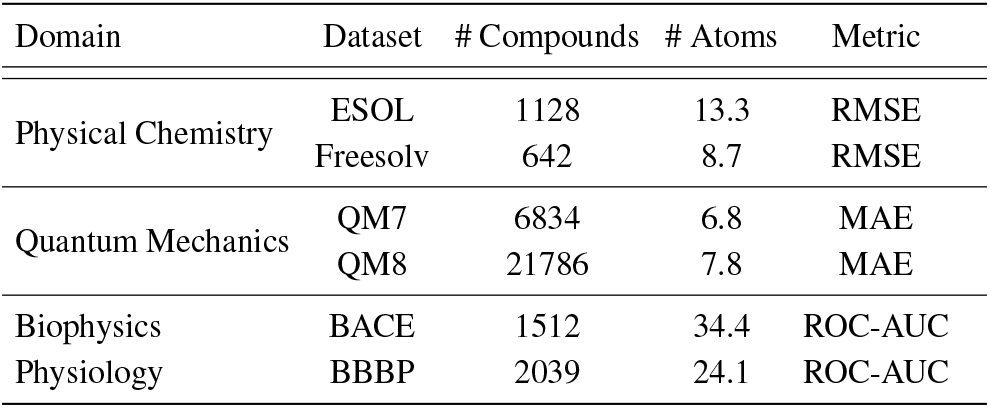
Summary of benchmark datasets. The table contains the applied domain, type, number of molecules, average number of atoms in the dataset, and metrics used.

- ESOL: Water solubility data (log solubility in mols per litre) for common organic small molecules.
- Freesolv Mobley and Guthrie (2014): Experimental and calculated hydration free energy of small molecules in water.
- QM7 Blum and Reymond (2009): Electronic properties (atomization energy, HOMO/LUMO, etc.) determined using ab-initio density functional theory (DFT).
- QM8 Ramakrishnan *et al*. (2015): Electronic spectra and excited state energy of small molecules calculated by multiple quantum mechanic methods.
- BBBP: Binary labels of blood-brain barrier penetration.
- BACE: Quantitative (IC50) and qualitative (binary label) binding results for a set of inhibitors of human *β*-secretase 1(BACE-1).

We select the six molecule datasets from MoleculeNet (Wu *et al*., 2018). The first two, ESOL and Freesolv, are physical chemistry datasets, and the next two, QM7 and QM8, are quantum mechanic datasets, then BACE and BBBP are biophysics and physiology, respectively. We also add 3D locational information to downstream datasets in the same way (Fang *et al*., 2022) as the pre-training dataset.

### 4.2 Training and evaluation settings

#### 4.2.1 Pre-training Setting

During the pre-training of the model, we use an Adam optimizer with an initial learning rate of 0.001 and exponentially reduce the learning rate at a ratio of 0.95 or 0.99. Pre-training process is run with 300 epochs and 400 batch sizes.

#### 4.2.2 Fine-tuning Setting

After pre-training, we fine-tuned the model on six benchmark datasets. We train the model with an Adam optimizer of a learning rate of 0.001 and exponentially decrease the learning rate at rates of 0.95 or 0.99 in the same way as pre-training. We set 32 batch sizes and ran 200 epochs for all datasets.

We split the data set into train/validation/test sets at a ratio of 80/10/10 using the scaffold splitter (Bemis and Murcko, 1996) from DeepChem (Ramsundar *et al*., 2019) for downstream tasks like previous works (Hu *et al*., 2019; Rong *et al*., 2020). Each set is structurally different as the scaffold splitter splits molecular data by their substructure. Then, this splitting method is widely used for molecular-related tasks because it can evaluate the generalization capability of algorithms well. We provide further details of hyperparameter settings in supplemental materials.

#### 4.2.3 Implementation Details

We implement our 3DGCL with PyTorch (Paszke *et al*., 2019) and PyTorch Geometric frameworks (Fey and Lenssen, 2019) and write our implementation code based on (Liu *et al*., 2021a). We use RDKit (Landrum *et al*., 2020), a cheminformatics software for molecular-related tasks. All the experiments are run on a single NVIDIA RTX 3090 GPU.

There are two processes regarding the running time of tasks about conformers: creating a conformer pool from the original ESOL dataset and contrastive learning after selecting two conformers. In our study, each process took 3 and 6 minutes, totaling approximately 9 minutes, and the number of model parameters is about 0.5 million. This indicates that our work requires significantly low resources.

#### 4.2.4 Evaluation

To evaluate our fine-tuned model, we measure the RMSE (Root Mean Squared Error) on ESOL, Freesolv, and the MAE (Mean Absolute Error) on QM7 and QM8 datasets. For a fair comparison with the state-of-the-art models, we run the model with random seeds three times and average the performance and standard deviation in the same way in previous works. In the same way (Wang *et al*., 2022; Rong *et al*., 2020; Zhou *et al*., 2022), we evaluated QM7 for 1 target task and QM8 for the average of 12 target tasks.

### 4.3 Results

We evaluate the 3DGCL performance with standard supervised baselines and self-supervised models. All compared methods use more than one dataset of six benchmark datasets (ESOL, Freesolv, QM7, QM8, BBBP, and BACE) and conduct experiments under the same condition (Zhou *et al*., 2022). The same condition denotes scaffold-splitting (with considering chirality) the train/validation/test data to an 8:1:1 ratio and running tests independently three times with three random seeds. Scaffold splitting can be divided into two types according to the consideration of chirality. We show the results of scaffold splitting with considering chirality in Table 3, the results without considering chirality in Supplementary Table S2.

**Table 3.** Test performance of 3DGCL and other methods based on four regression (ESOL, Freesolv, QM7, and QM8) and two classification benchmarks (BBBP and BACE). We mark the best results in bold and the second best results in <u>underlined</u>. We split the dataset into 8:1:1 (train:validation:test) using scaffold splitting considering chirality.

| Metric<br>Model | RMSE (Lower is better) ↓ |  | MAE (Lower is better) ↓ |  | ROC-AUC (Higher is better) ↑ |  |
| --- | --- | --- | --- | --- | --- | --- |
|  | ESOL | Freesolv | QM7 | QM8 | BBBP | BACE |
| DMPNN | 1.050 (0.008) | 2.082 (0.082) | 103.5 (8.6) | 0.0190 (0.0001) | 71.0 (0.3) | 80.9 (0.6) |
| Attentive FP | 0.877 (0.029) | 2.073 (0.183) | 72.0 (2.7) | 0.0179 (0.0001) | 64.3 (1.8) | 78.4 (0.02) |
| HMGNN | 0.832 (0.010) | 1.857 (0.071) | 59.0 (3.4) | 0.0173 (0.004) | 64.3 (1.8) | 78.4 (0.02) |
| N-Gram<sub>RF</sub> | 1.074 (0.107) | 2.688 (0.085) | 92.8 (4.0) | 0.0236 (0.0006) | 69.7 (0.6) | 77.9 (1.5) |
| N-Gram<sub>XGB</sub> | 1.083 (0.082) | 5.061 (0.744) | 81.9 (1.9) | 0.0215 (0.0005) | 69.1 (0.8) | 79.1 (1.3) |
| PretrainGNN | 1.100 (0.006) | 2.764 (0.002) | 113.2 (0.6) | 0.0200 (0.0001) | 68.7 (1.3) | 84.5 (0.7) |
| MolCLR | 1.271 (0.033) | 2.594 (0.249) | 66.8 (2.3) | 0.0178 (0.0003) | 72.2 (2.1) | 82.4 (0.9) |
| GraphMVP | 1.029 (0.033) | — | — | — | 72.4 (1.6) | 81.2 (0.9) |
| GROVER<sub>base</sub> | 0.983 (0.090) | 2.176 (0.052) | 94.5 (3.8) | 0.0218 (0.0004) | 70.0 (0.1) | 82.6 (0.7) |
| GROVER<sub>large</sub> | 0.895 (0.017) | 2.272 (0.051) | 92.0 (0.9) | 0.0224 (0.0003) | 69.5 (0.1) | 81.0 (1.4) |
| GEM | 0.798 (0.029) | 1.877 (0.094) | 58.9 (0.8) | 0.0171 (0.0001) | 72.4 (0.4) | <u>85.6</u> (0.2) |
| Uni-Mol | <u>0.788</u> (0.029) | <u>1.620</u> (0.035) | **41.8** (0.2) | <u>0.0156</u> (0.0001) | <u>72.9</u> (0.6) | **85.7** (0.2) |
| **3DGCL** | **0.778** (0.102) | **1.441** (0.19) | <u>42.53</u> (7.69) | **0.0143** (0.0001) | **79.15** (0.04) | 85.5 (0.03) |

The baselines are as follows: DMPNN (Yang *et al*., 2019b) proposes an interactive message passing scheme considering the interactions. AttentiveFP (Xiong *et al*., 2019) is an attention-based graph neural network. HMGNN (Shui and Karypis, 2020) utilizes global molecular representations using an attention mechanism. CD-MVGNN (Ma *et al*., 2022) performs a cross-dependent message-passing scheme considering both atom and bond information. We compare CD-MVGNN with the baselines in Supplementary Table S2 for the same test settings.

The rest of the seven methods are self-supervised models. N-Gram (Liu *et al*., 2019) produces a graph representation in short walks by building the node embedding. MolCLR (Wang *et al*., 2022) is a 2D-2D view contrastive learning model based on atom masking, bond deletion, and subgraph removal. GraphMVP (Liu *et al*., 2021b) proposes 2D-3D view contrastive learning approaches. GROVER (Rong *et al*., 2020) uses a predictive pre-training strategy of motif-level. GEM Fang *et al*. (2022) and Uni-Mol design predictive self-supervised learning scheme using 3D molecular information. We reference the performance of the baselines in Uni-Mol (Zhou *et al*., 2022).

We present the experimental result with dataset size in pre-training to show the efficiency of our method, as can be seen in Table 3. We mark the best results in bold and underline the second best in Table 3. 3DGCL outperforms all other methods on four of six datasets and shows the second-best performance in QM7 and third-best in BACE. Furthermore, our method achieves overwhelming performance on Freesolv and BBBP by a large margin.

The results present that 3DGCL consistently achieves the best performance and comparable results. We should also note that our method uses overwhelmingly fewer resources than other methods, as shown in Figure 3. We compare 3DGCL and the three state-of-the-art, GROVER, GEM, and Uni-Mol, that show the best overall performance with the pre-train dataset and model parameters. We can see that 3DGCL uses about 10,000 times fewer datasets using only 1k than other state-of-the-art methods using over 10M datasets, as shown in Figure 3(a). We can also confirm that the number of 3DGCL parameters is approximately over 100 times fewer than the parameter size of other models, as shown in Figure 3(b).

**Fig. 3.**
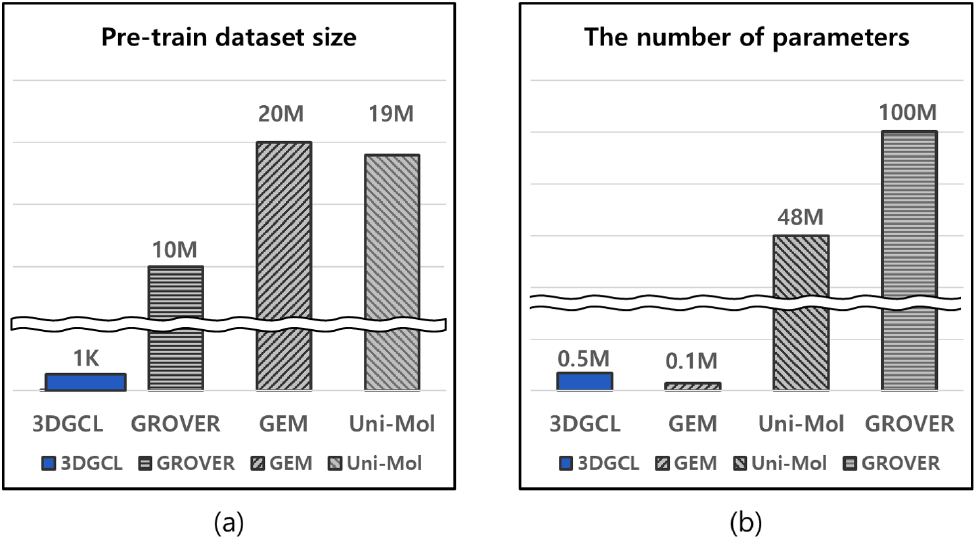
Comparison of 3DGCL and the three state-of-the-art models in terms of dataset and parameter size in pre-training.

## 5 Additional Experiments

### 5.1 Pre-training effects according to the molecular 3D information and size

3D information is molecular coordinates, and we can obtain geometric information such as distance, angle, and torsion. Our encoder utilize distance information in the pre-training process. To verify the contribution of geometric information on molecular representation learning, we compare our model with pre-training and the model without pre-training, i.e., SchNet. As a result of the experiment, 3DGCL consistently obtains enhanced performance on all the datasets, as shown in Table 4.

**Table 4.** Performance improvements of 3DGCL pre-training. We show enhanced performances to verify the pre-training effect of our method. We also indicate the number of atoms of datasets.

| Dataset<br>Molecular size | ESOL<br>13.3 | Freesolv<br>8.7 | QM7<br>6.8 | QM8<br>7.8 | BBBP<br>34.4 | BACE<br>24.1 |
| --- | --- | --- | --- | --- | --- | --- |
| without pre-training | 1.031 (0.1) | 1.91 (0.42) | 59.70 (22.1) | 0.0163 (0.0003) | 71.19 (0.03) | 76.2 (0.02) |
| with 3DGCL pre-training | 0.778 (0.102) | 1.441 (0.19) | 42.53 (7.69) | 0.0143 (0.0001) | 79.15 (0.04) | 85.5 (0.03) |
| **Improvement** | **24.5%** | **24.61%** | **28.8%** | **12.3%** | **10.1%** | **11%** |

We also investigate the pre-training effect by molecular size. The size denotes the average number of atoms in each dataset. The result shows that 3DGCL leads to higher self-supervised performance on small molecules than on larger datasets. This confirms the effect of the backbone encoder (SchNet), which was developed to focus on small organic molecules. We demonstrate that 3D spatial information is critical for molecular property prediction, and the encoder plays an important role in self-supervised learning.

### 5.2 Comparison with diverse augmentation methods

We conduct comprehensive studies to compare the proposed 3DGCL method with the existing contrastive learning approaches, which may alter the original molecular properties. In addition, we also compare our conformer pool, which consists of five conformers that can most likely exist in nature, and molecules that are difficult to exist in nature, which is created by adding noise to the 3D position of the original molecule. Finally, we compared selecting the most stable molecules with our method of selecting molecules at random. We visualize the methods used in this study in Figure 4.

**Fig. 4.**
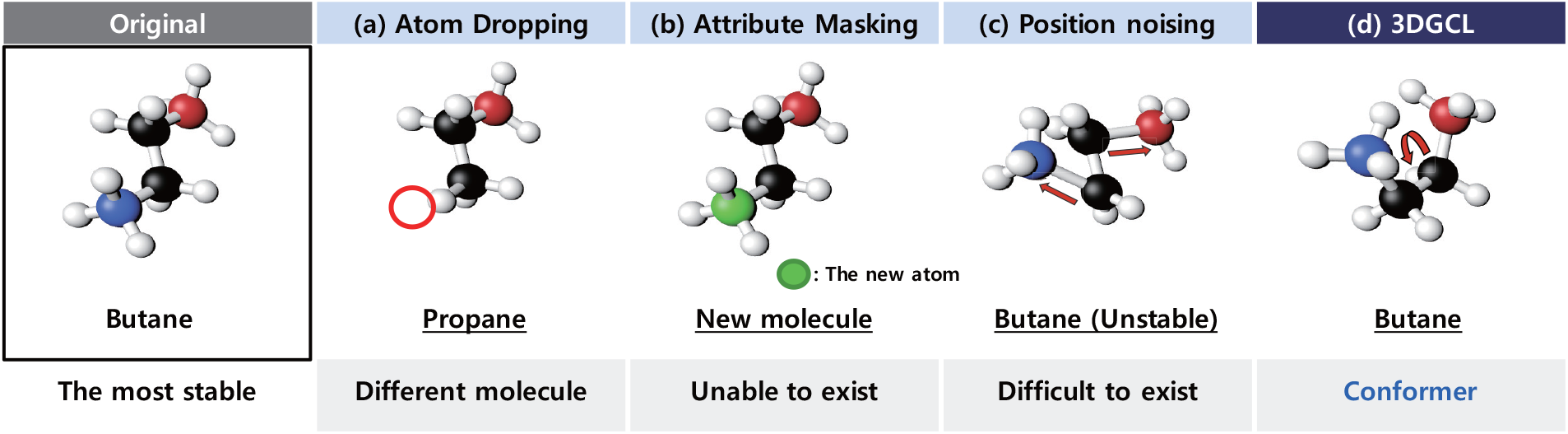
The overview of various molecular augmentation methods.

#### Atom Dropping

Atom dropping removes atoms randomly at a constant ratio in the molecule graph 𝒢. In previous graph contrastive learning studies, atom dropping has been widely used for graph augmentation (You *et al*., 2020; Wang *et al*., 2022). We randomly delete 20% of the atoms in the original molecule.

#### Attribute Masking

Like atom dropping, we randomly select 20% of atoms in the original molecule, then mask their attributes with a feature not presented in the original molecules (You *et al*., 2020; Wang *et al*., 2022). Since we use only the atomic number as the feature of atoms, we replace the atom number with the new masking number *n*.

#### Position Noising

We add noise to the original 3D coordinates of the most stable molecule. We generate the noise drawn from a Gaussian distribution ∼ 𝒩 (0, 1) whose mean and standard deviation are 0 and 1, then multiply the noise by 0.01 to fit the scale with coordinates.

#### Conformer-S

We compare the method using the two most stable conformers without creating the conformer pool to verify that the proposed way of randomly selecting in the conformer pool is effective for learning rich molecular representations. We name the approach utilizing only the two stable molecules conformer-S and call our method **Conformer-R** in this experiment.

We conduct extensive experiments to confirm the efficacy of molecular location information based on chemical knowledge. The experiments consist of three concepts. First, we compare atom dropping and atom masking methods, which are challenging to preserve molecular semantics due to severe structural changes. Secondly, we performed pre-training using the noising position method that utilizes 3D information while subtly giving spatial changes without chemical knowledge. Finally, we witness that our random select way is superior to the stable molecule choice method.

As experimental results, attribute masking has the least improved effect in pre-training, and atom dropping shows the second minor effect after attribute masking, as shown in Table 5. These results show that the method of significantly modifying the structure and properties of the original molecule is not suitable for molecular representation learning. Augmentation, position noising, which generates molecules that are difficult to exist (finely changing the 3D locations of atoms at random), showed a better effect than the above two approaches. We also test comparing the conformer pool (Conformer-R) and without the pool (Conformer-S) to confirm that our method plentifully learns from molecular geometric representation. As a result, our 3DGCL shows the best performance than other methods. Through diverse comparative experiments, we validate that our proposed approach is the most proper for molecular graph contrastive learning in the context of molecular property prediction.

**Table 5.** Details of experimental results based on diverse augmentation methods. We conduct extensive tests to prove the superiority of our conformer-based augmentation method for molecular property prediction. The methods are categorized according to semantic preservation, domain knowledge, and 3D information, and their results are reported.

| Method | ESOL | Freesolv | QM7 | QM8 | BBBP | BACE | Maintaining semantics | Chemical knowledge | 3D information |
| --- | --- | --- | --- | --- | --- | --- | --- | --- | --- |
| Atom Dropping | 0.808 | 1.743 | 57.14 | 0.0164 | 71.68 | 84.61 | X | X | X |
| Attribute Masking | 0.800 | 1.817 | 48.75 | 0.0171 | 75.17 | 84.81 | X | X | X |
| Position Noising | 0.780 | 1.89 | 51.34 | 0.0166 | 77.53 | 84.40 | X | X | O |
| Conformer-S | 0.787 | 1.743 | 48.73 | 0.0155 | 75.82 | 85.34 | O | O | O |
| **Conformer-R (Ours)** | **0.778** | **1.441** | **42.53** | **0.0143** | **79.15** | **85.52** | **O** | **O** | **O** |

### 5.3 Application to real world

The drug field suffers from annotation scarcity in that wet-lab experiments are required. We design experiments to verify that our model can be applied to the actual drug area, assuming the real world with a data deficiency. We conduct the experiments using ESOL and only 150, 300, and 600 samples as training data without using the 1,128 entire datasets.

Then, we compare the model’s results with pre-training and the model without pre-training. The pre-training process was conducted using the whole ESOL dataset in the same way as in the above experiments. We evaluate the results based on RMSE and observe that our method consistently outperforms the model trained from scratch in all the reduced datasets, as shown in Figure 5. The results indicate that our approach can be utilized in the drug field encountering annotation insufficiency to obtain significant effects.

**Fig. 5.**
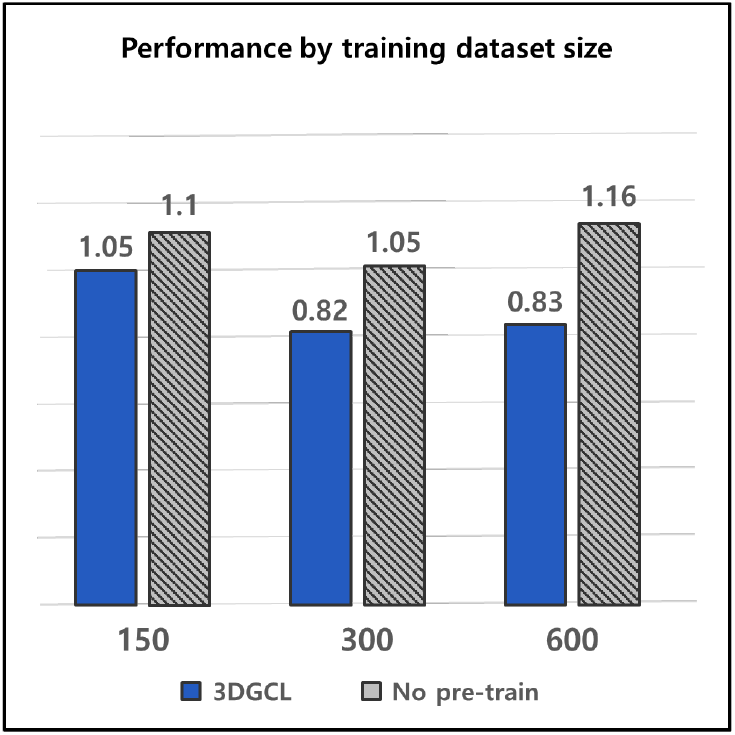
Comparision of performance difference by training dataset size.

## 6 Conclusion and Futher Works

In this work, we first proposed a novel 3D-3D view Graph Contrastive Learning (3DGCL) framework to address existing issues in self-supervised learning for molecular property prediction. Despite the prevailing notion that self-supervised learning requires hyper-scale resources, we present the possibility of small-scale self-supervised learning methods. Furthermore, we suggested a contrastive approach using randomization in a conformer pool that can learn 3D information in abundance while maintaining molecular semantics, unlike previous methods that could alter the property of the molecule. Comprehensive experiments show that our method outperforms existing self-supervised methods. We provide insight that it is proper to leverage 3D geometric information and domain-based knowledge in molecular property prediction. We also demonstrate that 3DGCL can be applied to the actual drug field undergoing annotation scarcity. We have shown remarkable performance using models and datasets, even on a small scale.

Despite the strong strengths, our model 3DGCL also has some weaknesses, such as focusing on only distance-based geometric information due to our 3D-based backbone framework. Another weakness is that the model needs to generate 3D information of molecules, which can slow the model training. If sufficient resources are supported, we will extend 3DGCL to further studies with a broader range of downstream tasks, along with hyper-scale model capacity, and more diverse geometric information.

## References

Adams, K., Pattanaik, L., and Coley, C. W. (2021). Learning 3d representations of molecular chirality with invariance to bond rotations. arXiv preprint arXiv:2110.04383.

Bemis, G. W. and Murcko, M. A. (1996). The properties of known drugs. 1. molecular frameworks. Journal of medicinal chemistry, 39(15), 2887–2893.

Blum, L. C. and Reymond, J.-L. (2009). 970 million druglike small molecules for virtual screening in the chemical universe database gdb-13. Journal of the American Chemical Society, 131(25), 8732–8733.

Chen, T., Kornblith, S., Norouzi, M., and Hinton, G. (2020). A simple framework for contrastive learning of visual representations. In International conference on machine learning, pages 1597–1607. PMLR.

Chithrananda, S., Grand, G., and Ramsundar, B. (2020). Chemberta: Large-scale self-supervised pretraining for molecular property prediction. arXiv preprint arXiv:2010.09885.

Danel, T., Spurek, P., Tabor, J., Śmieja, M., Struski, Ł., Słowik, A., and Maziarka, Ł. (2020). Spatial graph convolutional networks. In International Conference on Neural Information Processing, pages 668–675. Springer.

Delaney, J. S. (2004). Esol: estimating aqueous solubility directly from molecular structure. Journal of chemical information and computer sciences, 44(3), 1000–1005.

Devlin, J., Chang, M.-W., Lee, K., and Toutanova, K. (2018). Bert: Pretraining of deep bidirectional transformers for language understanding. arXiv preprint arXiv:1810.04805.

Dillard, L. (2021). Self-supervised learning for molecular property prediction.

Fang, X., Liu, L., Lei, J., He, D., Zhang, S., Zhou, J., Wang, F., Wu, H., and Wang, H. (2022). Geometry-enhanced molecular representation learning for property prediction. Nature Machine Intelligence, 4(2), 127–134.

Fey, M. and Lenssen, J. E. (2019). Fast graph representation learning with pytorch geometric. arXiv preprint arXiv:1903.02428.

Ganea, O., Pattanaik, L., Coley, C., Barzilay, R., Jensen, K., Green, W., and Jaakkola, T. (2021). Geomol: Torsional geometric generation of molecular 3d conformer ensembles. Advances in Neural Information Processing Systems, 34, 13757–13769.

Gilmer, J., Schoenholz, S. S., Riley, P. F., Vinyals, O., and Dahl, G. E. (2017). Neural message passing for quantum chemistry. In International conference on machine learning, pages 1263–1272. PMLR.

Halgren, T. A. (1996). Merck molecular force field. i. basis, form, scope, parameterization, and performance of mmff94. Journal of computational chemistry, 17(5-6), 490–519.

He, K., Fan, H., Wu, Y., Xie, S., and Girshick, R. (2020). Momentum contrast for unsupervised visual representation learning. In Proceedings of the IEEE/CVF conference on computer vision and pattern recognition, pages 9729–9738.

Hermosilla, P. and Ropinski, T. (2022). Contrastive representation learning for 3d protein structures. arXiv preprint arXiv:2205.15675.

Hu, W., Liu, B., Gomes, J., Zitnik, M., Liang, P., Pande, V., and Leskovec, J. (2019). Strategies for pre-training graph neural networks. arXiv preprint arXiv:1905.12265.

Klicpera, J., Groß, J., and Günnemann, S. (2020). Directional message passing for molecular graphs. arXiv preprint arXiv:2003.03123.

Landrum, G., Tosco, P., Kelley, B., sriniker, gedeck, NadineSchneider Vianello, R., Ric Dalke, A., Cole, B., AlexanderSavelyev, Swain, M., Turk, S., N, D., Vaucher, A., Kawashima, E., Wójcikowski, M., Probst, D., guillaume godin, Cosgrove, D., Pahl, A. JP, Berenger, F., strets123, JLVarjo, O’Boyle, N., Fuller, P., Jensen, J. H., Sforna, G., and DoliathGavid (2020). rdkit/rdkit: 2020_03_1 (q1 2020) release.

Li, P., Wang, J., Qiao, Y., Chen, H., Yu, Y., Yao, X., Gao, P., Xie, G., and Song, S. (2021). An effective self-supervised framework for learning expressive molecular global representations to drug discovery. Briefings in Bioinformatics, 22(6), bbab109.

Liu, M., Luo, Y., Wang, L., Xie, Y., Yuan, H., Gui, S., Yu, H., Xu, Z., Zhang, J., Liu, Y., Yan, K., Liu, H., Fu, C., Oztekin, B. M., Zhang, X., and Ji, S. (2021a). DIG: A turnkey library for diving into graph deep learning research. Journal of Machine Learning Research, 22(240), 1–9.

Liu, S., Demirel, M. F., and Liang, Y. (2019). N-gram graph: Simple unsupervised representation for graphs, with applications to molecules. Advances in neural information processing systems, 32.

Liu, S., Wang, H., Liu, W., Lasenby, J., Guo, H., and Tang, J. (2021b). Pre-training molecular graph representation with 3d geometry. arXiv preprint arXiv:2110.07728.

Liu, Y., Wang, L., Liu, M., Zhang, X., Oztekin, B., and Ji, S. (2021c). Spherical message passing for 3d graph networks.

Lu, C., Liu, Q., Wang, C., Huang, Z., Lin, P., and He, L. (2019). Molecular property prediction: A multilevel quantum interactions modeling perspective. ArXiv, abs/1906.11081.

Ma, H., Bian, Y., Rong, Y., Huang, W., Xu, T., Xie, W., Ye, G., and Huang, J. (2022). Cross-dependent graph neural networks for molecular property prediction. Bioinformatics, 38(7), 2003–2009.

Mikolov, T., Chen, K., Corrado, G., and Dean, J. (2013). Efficient estimation of word representations in vector space. arXiv preprint arXiv:1301.3781.

Mobley, D. L. and Guthrie, J. P. (2014). Freesolv: a database of experimental and calculated hydration free energies, with input files. Journal of computer-aided molecular design, 28(7), 711–720.

Paszke, A., Gross, S., Massa, F., Lerer, A., Bradbury, J., Chanan, G., Killeen, T., Lin, Z., Gimelshein, N., Antiga, L., et al. (2019). Pytorch: An imperative style, high-performance deep learning library. Advances in neural information processing systems, 32.

Qiao, Z., Welborn, M., Anandkumar, A., Manby, F. R., and Miller, T. F. (2020). Orbnet: Deep learning for quantum chemistry using symmetry-adapted atomic-orbital features. The Journal of chemical physics, 153 12, 124111.

Ramakrishnan, R., Hartmann, M., Tapavicza, E., and Von Lilienfeld, O. A. (2015). Electronic spectra from tddft and machine learning in chemical space. The Journal of chemical physics, 143(8), 084111.

Ramsundar, B., Eastman, P., Walters, P., Pande, V., Leswing, K., and Wu, Z. (2019). Deep Learning for the Life Sciences. O’Reilly Media. https://www.amazon.com/Deep-Learning-Life-Sciences-Microscopy/dp/1492039837.

Rogers, D. and Hahn, M. (2010). Extended-connectivity fingerprints. Journal of Chemical Information and Modeling, 50(5), 742–754. PMID: 20426451.

Rong, Y., Bian, Y., Xu, T., Xie, W., Wei, Y., Huang, W., and Huang, J. (2020). Self-supervised graph transformer on large-scale molecular data. Advances in Neural Information Processing Systems, 33, 12559–12571

Schütt, K., Kindermans, P.-J., Sauceda Felix, H. E., Chmiela, S., Tkatchenko, A., and Müller, K.-R. (2017). Schnet: A continuous-filter convolutional neural network for modeling quantum interactions. Advances in neural information processing systems, 30.

Shi, C., Luo, S., Xu, M., and Tang, J. (2021). Learning gradient fields for molecular conformation generation. In International Conference on Machine Learning, pages 9558–9568. PMLR.

Shui, Z. and Karypis, G. (2020). Heterogeneous molecular graph neural networks for predicting molecule properties. In 2020 IEEE International Conference on Data Mining (ICDM), pages 492–500. IEEE.

Stärk, H., Beaini, D., Corso, G., Tossou, P., Dallago, C., Günnemann, S., and Liò, P. (2021). 3d infomax improves gnns for molecular property prediction. arXiv preprint arXiv:2110.04126.

Sun, M., Xing, J., Wang, H., Chen, B., and Zhou, J. (2021). Mocl: Contrastive learning on molecular graphs with multi-level domain knowledge. ArXiv, abs/2106.04509.

Unke, O. T. and Meuwly, M. (2019). Physnet: A neural network for predicting energies, forces, dipole moments, and partial charges. Journal of Chemical Theory and Computation, 15(6), 3678–3693. PMID: 31042390.

Vaswani, A., Shazeer, N., Parmar, N., Uszkoreit, J., Jones, L., Gomez, A. N., Kaiser, Ł., and Polosukhin, I. (2017). Attention is all you need. Advances in neural information processing systems, 30.

Wang, S., Guo, Y., Wang, Y., Sun, H., and Huang, J. (2019). Smiles-bert: large scale unsupervised pre-training for molecular property prediction. In Proceedings of the 10th ACM international conference on bioinformatics, computational biology and health informatics, pages 429–436.

Wang, Y., Wang, J., Cao, Z., and Barati Farimani, A. (2022). Molecular contrastive learning of representations via graph neural networks. Nature Machine Intelligence, 4(3), 279–287.

Weininger, D. (1988). Smiles, a chemical language and information system. 1. introduction to methodology and encoding rules. Journal of chemical information and computer sciences, 28(1), 31–36.

Wu, Z., Ramsundar, B., Feinberg, E. N., Gomes, J., Geniesse, C., Pappu, A. S., Leswing, K., and Pande, V. (2018). Moleculenet: a benchmark for molecular machine learning. Chemical science, 9(2), 513–530.

Xiong, Z., Wang, D., Liu, X., Zhong, F., Wan, X., Li, X., Li, Z., Luo, X., Chen, K., Jiang, H., et al. (2019). Pushing the boundaries of molecular representation for drug discovery with the graph attention mechanism. Journal of medicinal chemistry, 63(16), 8749–8760.

Yang, K., Swanson, K., Jin, W., Coley, C., Eiden, P., Gao, H., Guzman-Perez, A., Hopper, T., Kelley, B., Mathea, M., Palmer, A., Settels, V., Jaakkola, T., Jensen, K., and Barzilay, R. (2019a). Analyzing learned molecular representations for property prediction. Journal of Chemical Information and Modeling, 59(8), 3370–3388. PMID: 31361484.

Yang, K., Swanson, K., Jin, W., Coley, C., Eiden, P., Gao, H., Guzman-Perez, A., Hopper, T., Kelley, B., Mathea, M., et al. (2019b). Analyzing learned molecular representations for property prediction. Journal of chemical information and modeling, 59(8), 3370–3388.

You, Y., Chen, T., Sui, Y., Chen, T., Wang, Z., and Shen, Y. (2020). Graph contrastive learning with augmentations. Advances in Neural Information Processing Systems, 33, 5812–5823.

Zhang, S., Hu, Z., Subramonian, A., and Sun, Y. (2020). Motif-driven contrastive learning of graph representations. arXiv preprint arXiv:2012.12533.

Zhou, G., Gao, Z., Ding, Q., Zheng, H., Xu, H., Wei, Z., Zhang, L., and Ke, G. (2022). Uni-mol: A universal 3d molecular representation learning framework.

Zhu, Y., Chen, D., Du, Y., Wang, Y., Liu, Q., and Wu, S. (2022). Featurizations matter: A multiview contrastive learning approach to molecular pretraining. In ICML 2022 2nd AI for Science Workshop.

