## Supplementary materials for "3D Graph Contrastive Learning for Molecular Property Prediction"

Table S1. Hyperparameter search for pre-training and fine-tuning. We select the best set of hyperparameters for our training processes.

| Hyperparemeter | Description | Range |
|---|---|---|
| **Pre-training settings** | | |
| batch size | the input batch size for the pre-training | 400 |
| learning rate | the initial learning rate for the pre-training | 0.001 |
| epochs | the number of epochs for the pre-training | 300 |
| z_dim | the hidden size for both encoder and projection head | [64, 128, 256] |
| activation | nonlinear function in projection head | ReLU |
| temperature ($\tau$) | the temperature parameter in NT-Xent loss | [0.2, 0.5] |
| gamma ($\gamma$) | exponential decay ratio of initial learning rate | [0.95, 0.99] |
| weight decay ($\lambda$) | L2 penalty for the pre-training | [0, 0.001] |
| dropout | dropout ratio for both encoder and projection head | 0.0 |
| **Encoder architecture** | | |
| cutoff | cutoff distance for interatomic interactions | [5.0, 8.0] |
| n_layers | the number of hidden layers in the encoder | 2 |
| filters | the number of filters in the encoder | [128, 256, 512] |
| gaussians | the number of gaussian functions | [50, 100] |
| **Fine-tuning settings** | | |
| batch size | the input batch size for the fine-tuning | 32 |
| learning rate | the initial learning rate for the fine-tuning | [0.001. 0.002] |
| epochs | the number of epochs for the fine-tuning | 200 |
| gamma ($\gamma$) | exponential decay ratio of initial learning rate | [0.95, 0.99] |
| weight decay ($\lambda$) | L2 penalty for the fine-tuning | [0, 0.001] |
| dropout | dropout ratio for fine-tuning | [0.0, 0.2, 0.3] |

### 1 Experimental setups

#### 1.1 Hyperparemeters settings

We run a random search to find the best combinations of hyperparameters for each downstream task. We select the hyperparameters with the best validation performance and report the test result during fine-tuning process. Table S1 shows all the hyperparameter lists we used in the experiments.

#### 1.2 Implementation Details

We implement our 3DGCL with PyTorch (Paszke *et al.*, 2019) and PyTorch Geometric frameworks (Fey and Lenssen, 2019) and write our implementation code based on (Liu *et al.*, 2021). We use RDKit (Landrum *et al.*, 2020), a cheminformatics software for molecular-related tasks. All the experiments are run on a single NVIDIA RTX 3090 GPU.

### 2 Dataset split

We select the molecule datasets from MoleculeNet (Wu *et al.*, 2018). We also add 3D conformer coordinates to datasets using the Merck molecular force field (MMFF94) (Halgren, 1996) function. Using scaffold splitting, we split the data set into train/validation/test sets at a ratio of 8/1/1. Scaffold split can lead to different results because the train/validation/test set composition differs depending on whether we consider chirality. We provide the additional results not considering chirality in Table S2.

### 3 Results

We evaluate the 3DGCL performance with standard supervised baselines and self-supervised models. All compared methods use more than one dataset of four benchmark datasets (ESOL, Freesolv, QM7, QM8) and conduct experiments under the same condition.

The baselines are as follows: GCN (Kipf and Welling, 2016) and Weave (Kearnes *et al.*, 2016) are graph convolution networks aggregating the neighbor node. As subsequent models of MPNN (Gilmer *et al.*, 2017), DMPNN (Yang *et al.*, 2019), MGCN (Lu *et al.*, 2019), and CoMPT (Chen *et al.*, 2021) propose an interactive message passing scheme considering the interactions. AttentiveFP (Xiong *et al.*, 2019) is an attention-based graph neural network. CD-MVGNN (Ma *et al.*, 2022) performs a cross-dependent message-passing scheme considering both atom and bond information.

The rest of the five methods are self-supervised models. MolCLR (Wang *et al.*, 2022) is a 2D-2D view contrastive learning model based on atom masking, bond deletion, and subgraph removal. 3Dinformax (Stärk *et al.*, 2021) proposes 2D-3D view contrastive learning approaches. GROVER (Rong *et al.*, 2020) uses a predictive pre-training strategy of motif-level. MPG Li *et al.* (2021) combines node and graph-level prediction tasks in pre-training. MEMO (Zhu *et al.*, 2022) is a multiple view contrastive learning method using various molecular representations (2D graph, 3D graph, Fingerprint, and SMILES). We reference the performance of the baselines in GROVER and their study.

We present the experimental result with dataset size in pre-training to show the efficiency of our method, as can be seen in Table S2. We mark the best results in bold and underline the second best in Table 3. The last column is the dataset size used in the pre-training process. 3DGCL outperforms all the baselines in the ESOL dataset and gains a 2.7% improvement over the second best model. In the Freesolv, MPG shows the best result, and 3DGCL gets the second great result. Our method obtains remarkable performance by a large margin in QM7.

### References

Chen, J., Zheng, S., Song, Y., Rao, J., and Yang, Y. (2021). Learning attributed graph representations with communicative message passing

Table S2. Test performance of 3DGCL and different methods based on four regression benchmarks. We mark the best results in bold and underline the second best. The last column is the dataset size used in the pre-training process. We split the dataset into 8:1:1 (train:validation:test) using scaffold splitting, not considering chirality. We reference the performance of the baselines in GROVER and their study.

| Metric | RMSE (Lower is better) ↓ | | MAE (Lower is better) ↓ | | |
|---|---|---|---|---|---|
| Model | ESOL | Freesolv | QM7 | QM8 | Pre-train Dataset |
| GCN | 1.211(0.052) | 3.174 (0.308) | 100.0 (3.8) | 0.0203 (0.0005) | |
| Weave | 1.158(0.055) | 2.398 (0.308) | 94.7 (2.7) | 0.022 (0.001) | |
| MPNN | 1.167(0.430) | 2.185 (0.952) | 113.0 (7.2) | 0.015 (0.002) | |
| DMPNN | 0.980(0.430) | 2.177 (0.914) | 105.8 (13.2) | 0.0143 (0.002) | |
| MGCN | 1.266(0.147) | 3.349 (0.097) | 77.6 (4.7) | 0.022 (0.002) | |
| AttentiveFP | 0.853(0.060) | 2.030 (0.420) | 126.7 (4.0) | 0.0282 (0.001) | |
| CoMPT | 0.878 (0.023) | 1.855 (0.578) | — | — | |
| CD-MVGNN | 0.779 (0.026) | 1.552 (0.123) | 70.358 (5.962) | **0.0124** (0.001) | |
| MolCLR | 1.11 (0.01) | 2.20(0.20) | 83.1 (4.0) | 0.0174 (0.0013) | 10,000k |
| 3Dinformax | 0.894 (0.028) | 2.337 (0.227) | — | — | 1,103k |
| MEMO | 0.984 (0.034) | — | — | — | 300k |
| GROVER | 0.831 (0.120) | 1.544 (0.397) | 72.5 (5.9) | 0.0125 (0.002) | 11,000k |
| MPG | 0.741 (0.017) | **1.269** (0.192) | — | — | 11,000k |
| **3DGCL** | **0.735 (0.08)** | 1.53 (0.42) | **41.33 (7.16)** | 0.0146 (0.0023) | **1k** |


transformer. *arXiv preprint arXiv:2107.08773*.

Fey, M. and Lenssen, J. E. (2019). Fast graph representation learning with pytorch geometric. *arXiv preprint arXiv:1903.02428*.

Gilmer, J., Schoenholz, S. S., Riley, P. F., Vinyals, O., and Dahl, G. E. (2017). Neural message passing for quantum chemistry. In *International conference on machine learning*, pages 1263–1272. PMLR.

Halgren, T. A. (1996). Merck molecular force field. i. basis, form, scope, parameterization, and performance of mmff94. *Journal of computational chemistry*, **17**(5-6), 490–519.

Kearnes, S., McCloskey, K., Berndl, M., Pande, V., and Riley, P. (2016). Molecular graph convolutions: moving beyond fingerprints. *Journal of computer-aided molecular design*, **30**(8), 595–608.

Kipf, T. N. and Welling, M. (2016). Semi-supervised classification with graph convolutional networks. *arXiv preprint arXiv:1609.02907*.

Landrum, G., Tosco, P., Kelley, B., sriniker, gedeck, NadineSchneider, Vianello, R., Ric, Dalke, A., Cole, B., AlexanderSavelyev, Swain, M., Turk, S., N, D., Vaucher, A., Kawashima, E., Wójcikowski, M., Probst, D., guillaume godin, Cosgrove, D., Pahl, A., JP, Berenger, F., strets123, JLVarjo, O'Boyle, N., Fuller, P., Jensen, J. H., Sforna, G., and DoliathGavid (2020). rdkit/rdkit: 2020_03_1 (q1 2020) release.

Li, P., Wang, J., Qiao, Y., Chen, H., Yu, Y., Yao, X., Gao, P., Xie, G., and Song, S. (2021). An effective self-supervised framework for learning expressive molecular global representations to drug discovery. *Briefings in Bioinformatics*, **22**(6), bbab109.

Liu, M., Luo, Y., Wang, L., Xie, Y., Yuan, H., Gui, S., Yu, H., Xu, Z., Zhang, J., Liu, Y., Yan, K., Liu, H., Fu, C., Oztekin, B. M., Zhang, X., and Ji, S. (2021). DIG: A turnkey library for diving into graph deep learning research. *Journal of Machine Learning Research*, **22**(240), 1–9.

Lu, C., Liu, Q., Wang, C., Huang, Z., Lin, P., and He, L. (2019). Molecular property prediction: A multilevel quantum interactions modeling perspective. In *Proceedings of the AAAI Conference on Artificial Intelligence*, volume 33, pages 1052–1060.
